# Structural basis of the Cope rearrangement and C–C bond-forming cascade in hapalindole/fischerindole biogenesis

**DOI:** 10.1101/173674

**Authors:** Sean A. Newmister, Shasha Li, Marc Garcia-Borràs, Jacob N. Sanders, Song Yang, Andrew N. Lowell, Fengan Yu, Janet L. Smith, Robert M. Williams, K. N. Houk, David H. Sherman

## Abstract

**STRUCTURES:** The atomic coordinates and structure factors for:

HpiC1 W73M/K132M SeMet (*P*2_1_2_1_2_1_) –1.7 Å

HpiC1 native (*C*2) –1.5 Å

HpiC1 native (*P*4_2_) –2.1 Å

HpiC1 Y101F (*C*2) –1.4 Å

HpiC1 Y101S (*C*2) –1.4 Å

HpiC1 F138S (*P*2_1_) –1.7 Å

HpiC1 Y101F/F138S (*P*2_1_ –1.65 Å
have been deposited with the Research Collaboratory for Structural Bioinformatics as Protein Data Bank entries 5WPP, 5WPR, 6AL6, 5WPR, 5WPU, 6AL7, and 6AL8 (www.rcsb.org).

**GRANTS:** This work was supported by: The authors thank the National Science Foundation under the CCI Center for Selective C-H Functionalization (CHE-1205646), the National Institutes of Health (CA70375 to RMW and DHS), R35 GM118101, R01 GM076477 and the Hans W. Vahlteich Professorship (to DHS) for financial support. M.G-B. thanks the Ramón Areces Foundation for a postdoctoral fellowship. J.N.S. acknowledges the support of the National Institute of General Medical Sciences of the National Institutes of Health under Award Number F32GM122218. Computational resources were provided by the UCLA Institute for Digital Research and Education (IDRE) and the Extreme Science and Engineering Discovery Environment (XSEDE), which is supported by the NSF (OCI-1053575). The content does not necessarily represent the official views of the National Institutes of Health.

**ABSTRACT:** Hapalindole alkaloids are a structurally diverse class of cyanobacterial natural products defined by their varied polycyclic ring systems and diverse biological activities. These polycyclic scaffolds are generated from a common biosynthetic intermediate by the Stig cyclases in three mechanistic steps, including a rare Cope-rearrangement, 6-*exo*-*trig* cyclization, and electrophilic aromatic substitution. Here we report the structure of HpiC1, a Stig cyclase that catalyzes the formation of 12-*epi*-hapalindole U in vitro. The 1.5 Å structure reveals a dimeric assembly with two calcium ions per monomer and the active sites located at the distal ends of the protein dimer. Mutational analysis and computational methods uncovered key residues for an acid catalyzed [3,3]-sigmatropic rearrangement and specific determinants that control the position of terminal electrophilic aromatic substitution leading to a switch from hapalindole to fischerindole alkaloids.

## INTRODUCTION

The hapalindole family of alkaloids are a large and structurally diverse class of natural products from aquatic and terrestrial cyanobacteria of the order *Stigonematales* (*1*). These metabolites are active against a broad range of biological targets, which include antibacterial, antifungal, insecticidal, and antimitotic activities (*2*-*7*). Each member is classified as a hapalindole, ambiguine, fischerindole, or welwitindolinone based on its core ring system (Fig. S1), and they have been the subject of various total syntheses due to their challenging structural complexity and unique biological properties (*1*). Until recently comparatively little was known regarding the biogenesis of these alkaloids, particularly with respect to the construction of the tetracyclic core ring system.

Initial reports demonstrated that the hapalindoles are derived from cis-indole isonitrile and geranyl pyrophosphate (GPP) (*8*, *9*), but the biogenesis of the polycyclic ring systems remained elusive. We recently identified an unexpected common biosynthetic intermediate **1** that undergoes a Cope rearrangement followed by a cyclization cascade to generate the hapalindole core (Fig. 1*a*) (*10*), offering a rare example of a pericyclic Cope rearrangement in natural product biosynthesis (*11*-*13*). This initial discovery was further expanded to include several Stig cyclases and revealed that the variant configurations observed in this class of alkaloids are generated from a central biosynthetic intermediate 1, transformed to products in a regio- and stereospecific fashion (Fig. 1*b*) (*14*-*16*). Thus, biogenesis of hapalindole-type metabolites includes a fascinating mechanistic puzzle regarding how homologous Stig cyclases mediate formation of varied hapalindoles or fischerindoles based on a remarkable C-C bond forming cascade. The biosynthesis of 12-*epi*-hapalindole U (**12H**) by HpiC1 is proposed to proceed through a three-part reaction mechanism: (*1*) Cope rearrangement of initial substrate **1** to generate intermediate **3**, which sets the stereochemistry at positions 11 and 12 in the final products; (*2*) 6-*exo*-*trig* cyclization of intermediate **3** to intermediate **4**, which sets the stereochemistry at positions 10 and 15 in the final products; and (*3*) electrophilic aromatic substitution of intermediate **4** to give **12H** upon deprotonation (Fig. 1*b*). This three-part reaction scheme is employed by the varied Stig cyclases which maintain exquisite stereochemical and regiochemical control through each of these mechanistic steps (*14*-*16*).

**Fig. 1.**
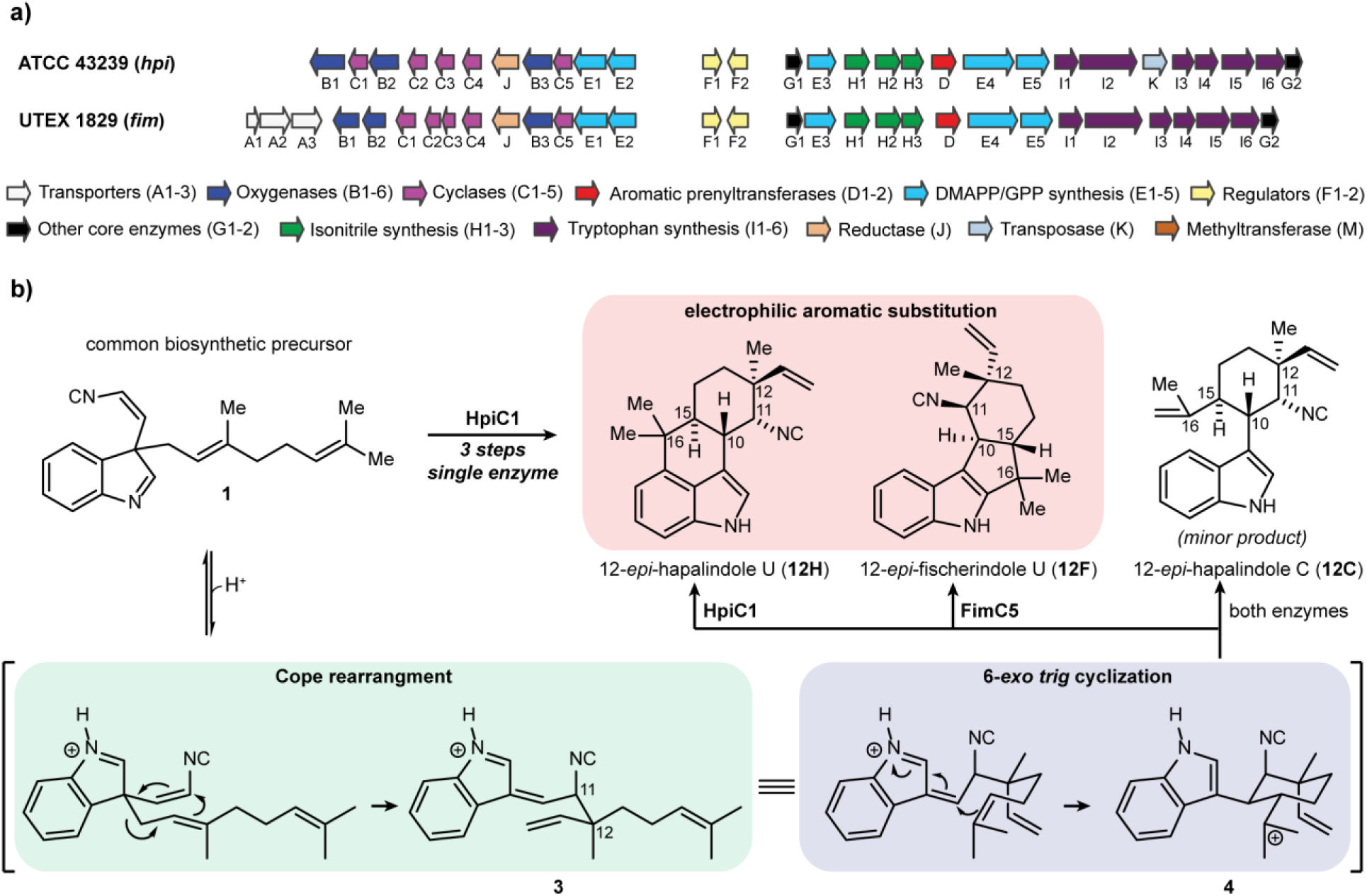
Biogenesis of hapalindole alkaloids. (**a**) *hpi* and *fim* gene clusters from *Stigonematales* cyanobacterial strains *Fischerella* sp. ATCC 43239 and *Fischerella muscicola* UTEX 1829. (**b**) The hapalindole and fischerindole core ring systems arise from the common biosynthetic intermediate **1**. Stig cyclases catalyze a Cope rearrangement followed by cyclization to generate tetracyclic products and trace levels of tri-cyclic shunt products. HpiC1 catalyzes formation of **12H**, while FimC5 catalyzes formation of **12F**, with identical stereochemistry at C11 and C12, but different C-ring regiochemistry.

In this study, we describe the molecular basis for the Stig cyclase ability to control three reactions, including an enzyme-catalyzed [3,3]-sigmatropic rearrangement, a 6-*exo*-*trig* cyclization, and a late-stage electrophilic aromatic substitution (Fig. 1). Originally annotated as unknown proteins (*8*), no previous information was available regarding the structure of the Stig cyclases. BLAST searches of the PDB database failed to identify matches, while models provided by the Phyre2 server (*17*) were based on carbohydrate binding modules of bacterial xylanases (*18*) that lack any homologous function to the Stig cyclase proteins.

We describe herein the first crystal structure of a Stig cyclase, and show through a mutational analysis that localizes the active site of HpiC1 (and other members of these unique enzymes) the ability to reconfigure its metabolite profile. These data combined with computational studies provide compelling insights into the mechanism of Cope rearrangement, 6-*exo*-*trig* cyclization and electrophilic aromatic substitution for this broad class of natural products. This work provides a strong foundation for understanding the selectivity of related cyclases that catalyze the formation of variant indole alkaloid metabolites while sharing common active site environments. These new findings will facilitate methods to engineer Stig cyclases and allied enzymes to achieve chemical diversification of biologically active hapalindole-type natural products.

## RESULTS

Following recent work that reported the function and selectivity of Stig cyclase proteins (*10*, *14*-*16*), we sought a crystal structure of HpiC1 in an effort to understand the structural basis of the complex cyclization cascade of its primary substrate **1**, and to understand how the Stig cyclases catalyze formation of variant alkaloid products from this common biosynthetic intermediate. Ultimately, we obtained HpiC1 crystals in four different forms, under conditions that differed in Ca^2+^ concentration. The initial 1.7 Å structure in Form 1 was solved by selenomethionyl SAD phasing from a HpiC1 W73M/K132M double mutant, as the wild-type protein lacks Met. The SeMet W73M/K132M structure was used as a search model to solve structures in the other three crystal forms by molecular replacement (Table S1). The overall fold of the HpiC1 polypeptide is a flattened β-jelly roll fold (Fig. 2*a*) composed of two antiparallel β-sheets. In all crystal forms, the antiparallel pairing of strands β6 in two monomers creates a continuous β-sheet across an extensive dimer interface, which buries 2060 Å^2^ of total surface area (PISA, Fig. 2*b*) (*19*), and encompasses approximately 20% of the total surface area of each monomer. This dimer interface is consistent with size exclusion chromatography analysis in which HpiC1 and other Stig cyclases migrated as apparent dimers (*14*). With the HpiC1 structure we conducted a DALI search, which showed highest homology (2.3 Å rmsd) with the carbohydrate-binding module (CBM) from xylanase in the thermostable bacterium *Rhodothermus marinus* (PDBid: 2Y64, Fig. 2*c*) (*20*). The proteins have highly similar tertiary structure and topology with the most significant differences occurring at their N-termini and in the loop regions between shared β-strands.

**Fig. 2.**
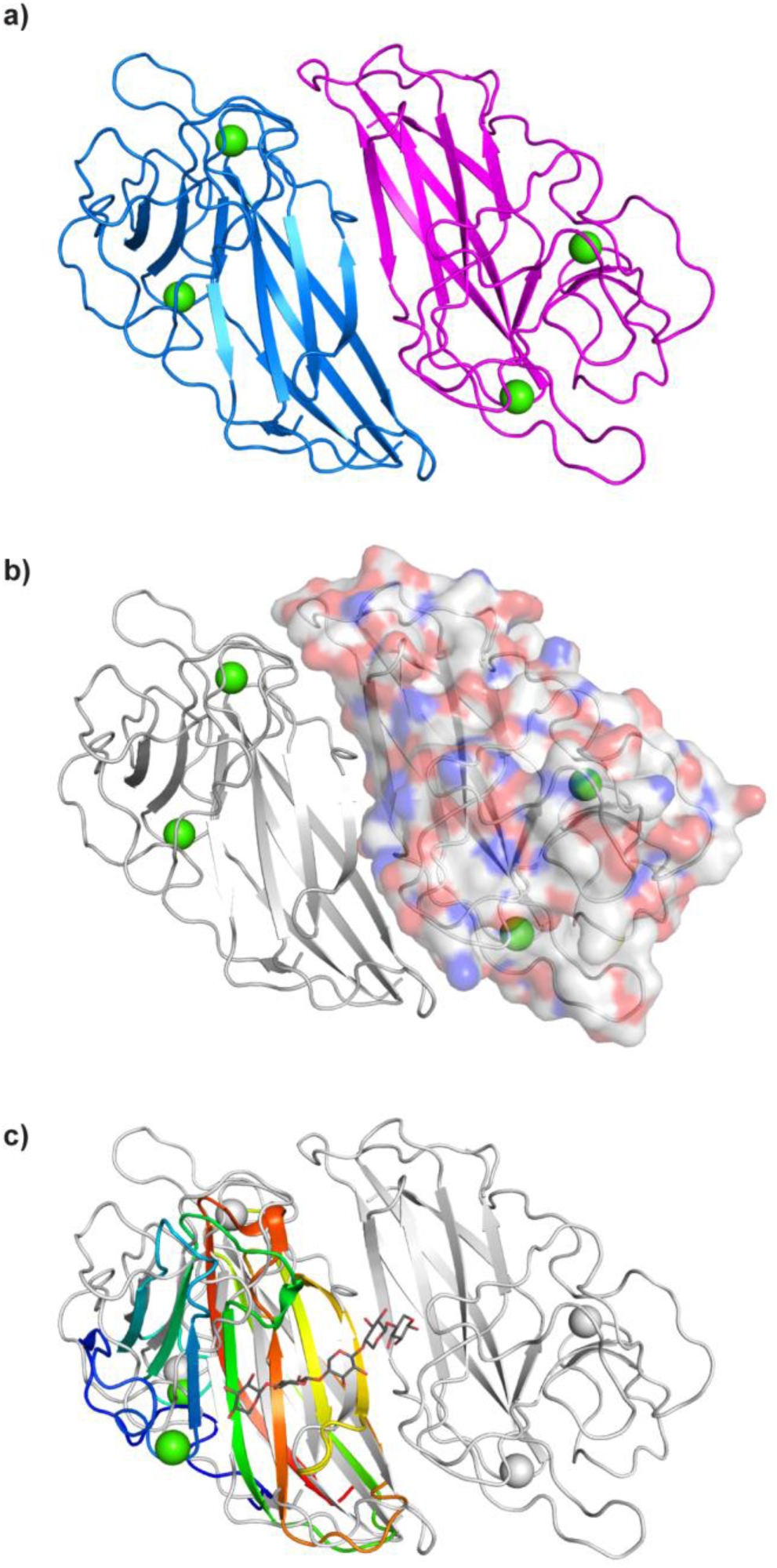
HpiC1 structure overview at 1.5 Å. (**a**) Cartoon representation of HpiC1 homodimer; the subunits are colored blue and magenta, green spheres indicate bound calcium ions. (**b**) Surface representation of a single protomer shows 2060 Å^2^ buried surface area between the subunits. (**c**) Superposition of xylanase CBM homolog (PDB id: 2Y64, rainbow); CBM is monomeric despite sharing the same fold as HpiC1.

HpiC1 has two integral Ca^2+^ ions, each with octahedral coordination geometry. Ca^2+^ site 1 is ligated by the carbonyl oxygens from Fl38, Ll47, Gl49, the side chain oxygens of Nl37 and Dl75, and a water molecule; and Ca^2+^ site 2 by the G37, E95 and N98 carbonyl oxygens and the E14, E95, and D216 side chains (Fig. S2). These sites were a key starting point to assess the structural and catalytic role that Ca^2+^ plays in the Stig cyclase enzymes. In vitro assays conducted in the presence of 5 mM EDTA showed no activity, leading us to conclude that Ca^2+^ is required for catalytic function in HpiC1 (*15*). From sequence comparisons, we expect the integral calcium-binding sites and the core dimeric assembly to be maintained in the other Stig cyclase enzymes (Fig. S3).

We next sought to determine the location of the enzyme active site. The high resolution, substrate-free HpiC1 structure did not immediately suggest a location for substrate binding and catalysis. The first indication of the enzyme active site location was revealed by abnormal difference density in a pocket located at the distal end of each subunit between Ca^2+^ sites 1 and 2. Polyethylene glycol from the crystallization was modeled into this pocket (Fig. S4), which was composed of numerous aromatic amino acids (Fig. 3*a*). Given the hydrophobicity of substrate **1**, we explored whether this hydrophobic pocket could indeed be the enzyme active site. This localized region was probed using Autodock VINA with **12H** (Fig. 1) as the ligand (*21*). The major product of the enzymatic reaction was chosen as the docking ligand based on its defined stereochemistry and rigid scaffold. The top three docking solutions had calculated affinities ranging from −9.7 – −9.4 kcal/mol and showed a fit and shape complementary with this hydrophobic pocket (Fig. S5), which suggested to a first approximation that this binding domain possesses an appropriate size and shape to accommodate the hapalindole core.

**Fig. 3.**
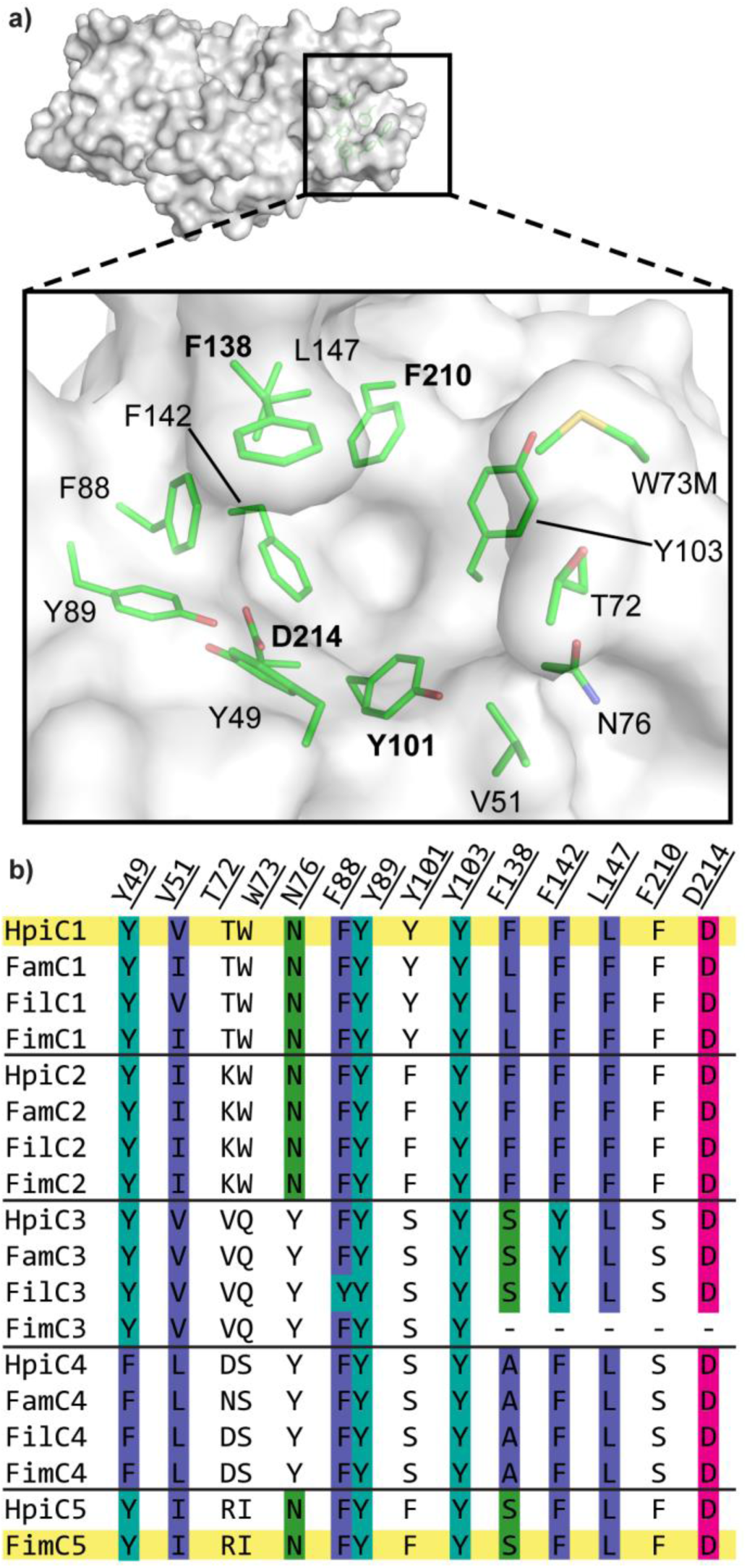
Active site of SeMet HpiC1 W73M/K132M. (**a**) Surface representation of the SeMet HpiC1 active site. Key residues are shown as green sticks. Met73 is substituted for the native tryptophan residue. This mutant protein retained native activity (**b**) Key active site residues shown in an alignment with other Stig cyclases. Residues are colored by conservation and side chain composition (ClustalX).

The initial docking solutions compelled us to further interrogate this region of the protein as the putative active site. First, we probed the role of HpiC1 D214 toward catalyzing formation of **12H**. This residue is 100% conserved in all of the currently identified Stig cyclases and is exceptional as the amino acid in the hydrophobic pocket most likely to participate in acid/base chemistry, which can reasonably be inferred to promote [3,3]-sigmatropic rearrangments (*22*). The HpiC1 structure shows that D214 carboxylate lacks a counter-ion and is hydrogen bonded to the Y89 hydroxyl. Substitution of D214 to alanine abolished activity in HpiC1 (Fig. 4), indicating that it plays a critical role in the catalytic cascade to generate cyclized products (addressed computationally below), and provides compelling evidence that this hydrophobic pocket is the enzyme active.

**Fig. 4.**
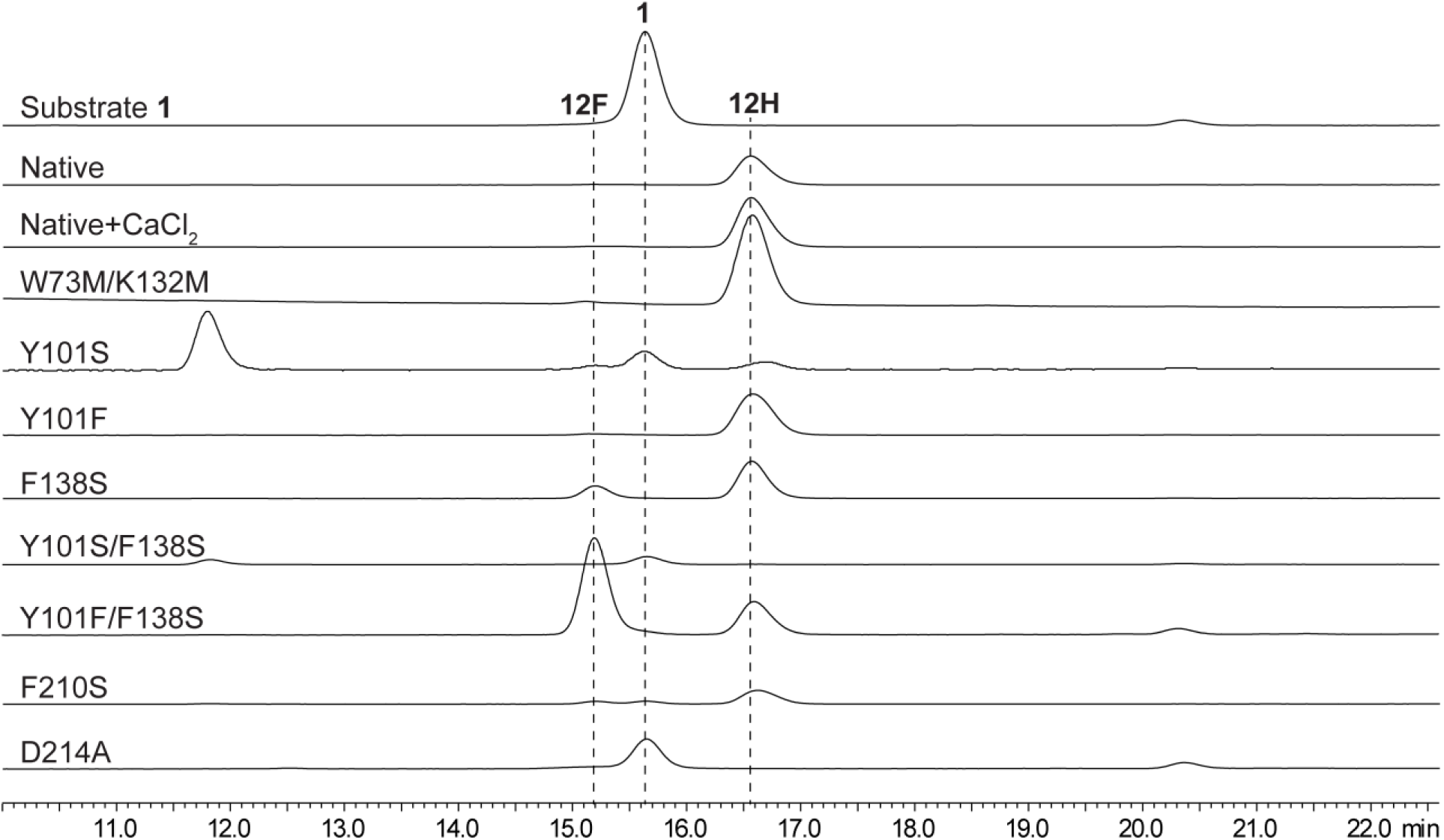
In vitro characterization of HpiC1 mutants using **1** as substrate. Substitution of the catalytic acid D214 to alanine abolished activity. A single mutation F138S altered the native product profile of HpiC1 to produce **12F**. The product became predominantly **12F** in the HpiC1 Y101F/F138S double mutant.

The enzyme conformation about D214 varies significantly for the HpiC1 structures solved among the individual crystal forms (Fig. S6). In Form 1, D214 is shielded by F138, whereas this residue is shifted in Forms 2, 3, and 4 to expose D214 to the binding pocket. The loop containing F138 is also shifted ~3 Å from its position in Form 1, although coordination of the structural calcium ion by the carbonyl oxygen of F138 is maintained. The protein atom B-factors in this region also suggest a higher degree of mobility at this site compared to other sections of the protein (Fig. S7), which is consistent with its role in recognition of an extended hydrocarbon substrate prior to cyclization. It is apparent that conformational flexibility would be required to perform multiple catalytic steps in a single binding domain, and also to accommodate the broad range of regio- and stereochemical configurations present in this class of polycyclic alkaloids derived from the initial bicyclic species **1** (*1*).

Based on the different product profiles of the various Stig cyclases, we examined the conservation of key residues in this hydrophobic pocket (Fig. 3*b*). Initial comparison of the Stig cyclases revealed that primary sequence clustering correlates to similar product profiles. We reasoned that information regarding key determinants of cyclase reactivity was contained within these localized sequences (Fig. 3), and a mutagenesis campaign was initiated to explore this active site. A comparative analysis between HpiC1 (catalyzes **12H** production from **1**) and FimC5 (catalyzes **12F** production from **1**) was initially pursued based on their function as homodimers in vitro, and their corresponding ability to differentially produce hapalindole or fischerindole core ring systems, respectively (Fig. 1) (*14*). Remarkably, substitution of F138 in HpiC1 to the corresponding serine from FimC5 led to the generation of a mixture of its major product, hapalindole **12H**, and the FimC5 major product, fischerindole **12F** (Fig. 4). The product ratio in F138S is approximately 1:1 (**12H**:**12F**) and is further shifted to 2:1 (**12F**:**12H**) in the Y101F/F138S double mutant. The corresponding mutation in FimC5 did not catalyze formation of **12H**; instead a shift in the product profile was observed in the FimC5 S139F mutant towards the production of **12C**, a minor tricyclic shunt product of the native HpiC1 and FimC5 reactions (Fig. S8) (*14*, *23*). These data (and computational studies described below) indicate that S139 plays a key role in directing terminal electrophilic aromatic substitution in FimC5.

Additional mutations were introduced into HpiC1 to identify key residues in the enzyme active site (Fig. 4), which were similarly guided by alignments with other Stig cyclases. Of particular interest was the Y101S/F138S HpiC1 variant that corresponds to the FamC3/HpiC3/FilC3 homologs. This mutant showed no activity with **1** in vitro, which is consistent with the inability of homodimeric forms of FamC3/HpiC3/FilC3 to generate products (*14*). Intriguingly, FamC3 was shown to associate with FamC2 as a heterodimer, and catalyze the formation of hapalindole H (*14*). The HpiC1 Y101S mutation also showed reduced activity, though to a lesser extent than in combination with F138S. HpiC1 Y101F had comparable activity to wild-type protein.

In order to further investigate the structural impact of these various mutations, high resolution structures were determined for HpiC1 variants Y101F, Y101S, F138S, and Y101F/F138S. While our efforts to observe bound ligands by either soaking or co-crystallization only afforded complexes with DMSO and Tris (Fig. S4), these structures significantly aided our efforts to interrogate the mechanism of cyclization in HpiC1 using computational methods. We first applied Molecular Dynamics (MD) simulations to the substrate-free structures to gain insights into the structure and dynamics of the enzyme active site and its impact on catalysis. Starting from the apo HpiC1 dimeric structure, analysis of the MD trajectories revealed large fluctuations of the loop containing F138 (N137-F150) (Fig. S9), in agreement with the different conformations found for this loop in the SeMet and native crystal structures (Fig. S6). The flexibility of the loop, which is in part controlled and reduced by a structural calcium ion, induces conformational changes in the F138 active site residue. These data show that F138 primarily explores two different conformations along the MD trajectory.

MD simulations show that D214, which is essential for enzyme activity as demonstrated by mutagenesis experiments described above, stays preferentially in a conformation in which the D214 side chain is pointing towards the inner cavity of the active site, while F138 acts as a wall on the side of the active site pocket (Fig. 5*a*, Fig. S10). This is due in part to the Y89 hydroxyl H-bond with the D214 carboxylic acid group that facilitates this arrangement (Fig. S10-S11). Importantly, the predicted pKa value of D214, estimated from different snapshots obtained along the 500ns of MD simulation (Fig. S12*a*) is 6.5 - 7.0, indicating that it can be protonated in an acid-base equilibrium to act as a protonating species during catalysis. Although the arrangement of the active site pre-organized for catalysis is the preferred one, an alternative conformation of F138 is sampled during the 500 ns trajectory. In this alternative conformation, the F138 side chain is displaced towards the center of the active site, and D214 becomes buried and inaccessible for interaction with the substrate, generating an inactive conformation (Fig. S10-S11). This alternative conformation is very similar to the active site arrangement observed in the SeMet crystal structure (Fig. S6).

**Fig. 5.**
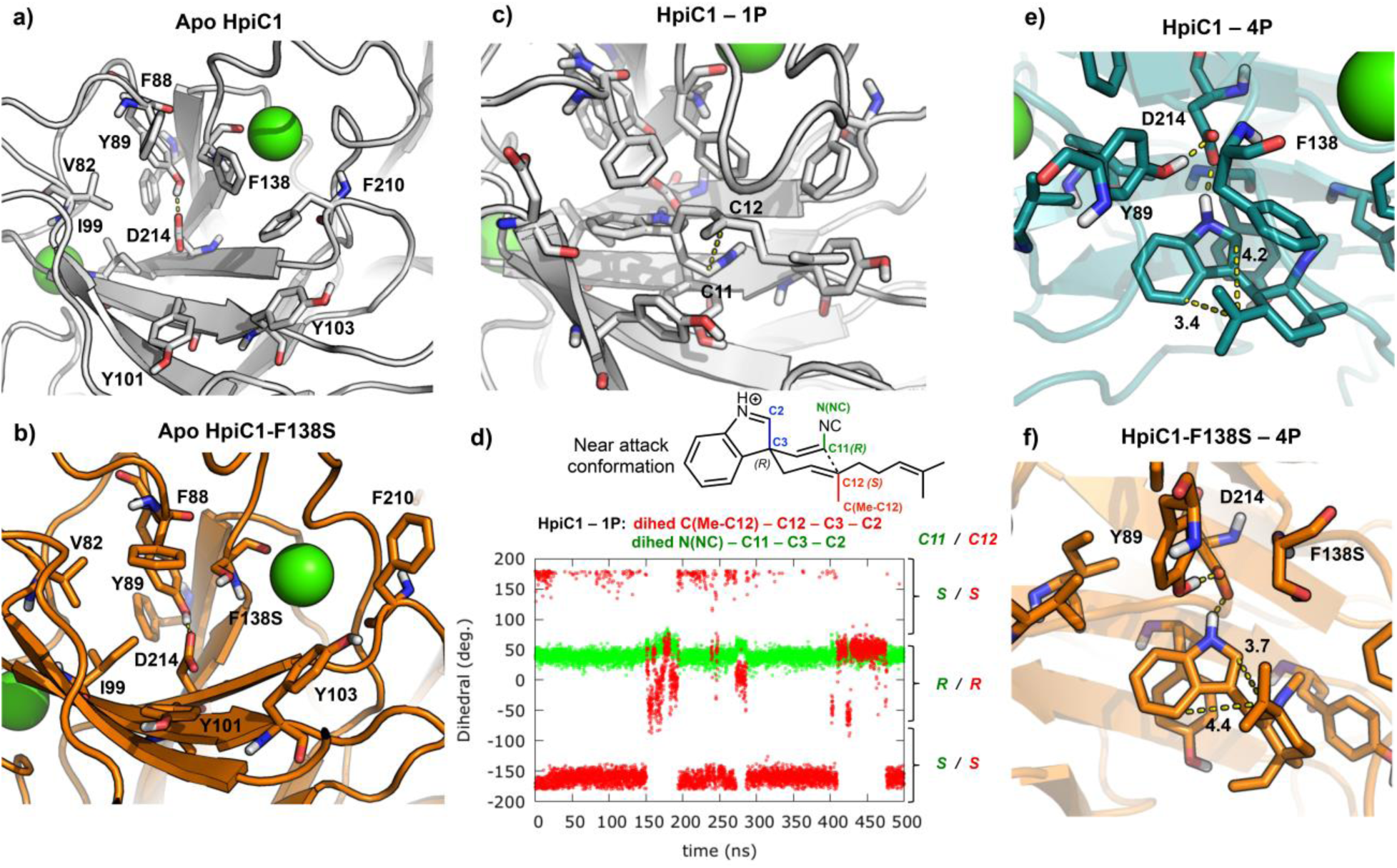
Representative snapshots obtained from 500 ns MD simulations of the active site arrangements in (**a**) apo HpiC1; (**b**) F138S single mutant; (**c**) substrate **1**(*R*)**P** bound into HpiC1; (**d**) Dihedral angles explored during the 500 ns of MD simulation for substrate **1**(*R*)**P** bound into HpiC1. Dihedral-1 (N(NC) – C11 – C3 – C2) and dihedral-2 (C(Me-C12) – C12 – C3 – C2) define the relative orientation of substituents at C11 and C12 positions, respectively, during the MD simulation. The right axis indicates the final stereochemistry of C11 and C12 expected after the Cope rearrangement coming from the given near attack conformation (NAC) of **1**(*R*)**P**, as shown in the scheme (**d**) and in Figure S21. **1**(*R*)**P** mainly explores one conformation during the MD trajectory leading to an (*R*) configuration at C11 and an (*S*) configuration at C12. Representative snapshots obtained from 500 ns MD simulations of the active site arrangements for intermediate **4P** bound into (**e**) HpiC1 (Fig. S22-S23); and (**f**) F138S single mutant (Fig. S24). Distances (in Å) show that the conformation adopted by intermediate **4P** in the HpiC1 wild-type enzyme moves C16 closer to C4 to form the hapalindole natural product, while the extra space in the F138S mutant enables the exploration of a new conformation of intermediate **4P** in which C16 gets closer to C2 to allow fischerindole product formation.

We next considered the absence of activity in the HpiC1 Y101S/F138S double mutant. MD simulations for Y101S/F138S in its apo state showed that the newly introduced serine Y101S interacts closely with the catalytic D214, in addition to the original Y89 hydroxyl H-bond (Fig. S13-S14). These two H-bonds between D214 and Y101S and Y89 favor stabilization of the negatively charged D214 carboxylate group, which will not be protonated in this more polar environment also favored by the neighboring F138S. This is further confirmed by the decrease of the predicted pKa value of D214 (pKa ≈ 5.5 - 6) (Fig. S12*b*). Moreover, D214 becomes more buried in the active site cavity, preventing its interaction with the substrate (Fig. S14).

As described above, F138S and Y101F/F138S mutants change the product profile in HpiC1 leading to increased formation of fischerindole **12F** (Fig. 4). We performed 500 ns MD simulations on both F138S and Y101F/F138S mutants, finding important changes in the active site pocket shape. The HpiC1 F138S mutation creates more space around the catalytic D214 residue (Fig. 5*b*) compared to wild-type enzyme containing the original F138 residue. This mutation also releases the interaction between the two phenyl rings of F138 and F210, with the F210 side chain becoming more flexible in HpiC1 F138S and Y101F/F138S mutants (Fig. 5*a*-b, Fig. S15). This active site reshaping is responsible for the change in the reaction outcome, as discussed below.

Next, we employed density functional theory (DFT) calculations to explore the possible reaction mechanism for Stig cyclases, in particular the [3,3]-sigmatropic rearrangement (Cope rearrangement), which is the first step in the HpiC1 catalyzed three-part reaction cascade starting from **1**. The instability of precursor **1** has precluded determining its chiral configuration at the indole C3 position. Thus, we computed the Cope rearrangement and cyclization cascade mechanism (Fig. 6) starting from (*R*)-**1**, the enantiomer predicted to be accepted by HpiC1 (as discussed below), and the corresponding mechanism starting from the (*S*)-**1** enantiomer (illustrated in Fig. S16). The biosynthesis of hapalindole **12H** by HpiC1 and fischerindole 12F by FimC5 (and related enzymes) is believed to proceed through three mechanistic steps: (1) the Cope rearrangement of initial substrate **1** to generate intermediate **3**, which sets the stereochemistry at positions 11 and 12 in the final products; (2) 6-*exo*-*trig* cyclization of intermediate **3** to intermediate **4**, which sets the stereochemistry at positions 10 and 15 in the final products; and (3) electrophilic aromatic substitution of intermediate **4** at two possible sites to give either intermediate **5H**, which yields **12H** upon deprotonation, or intermediate **5F**, which yields **12F** upon deprotonation (Fig. 6).

**Fig. 6.**
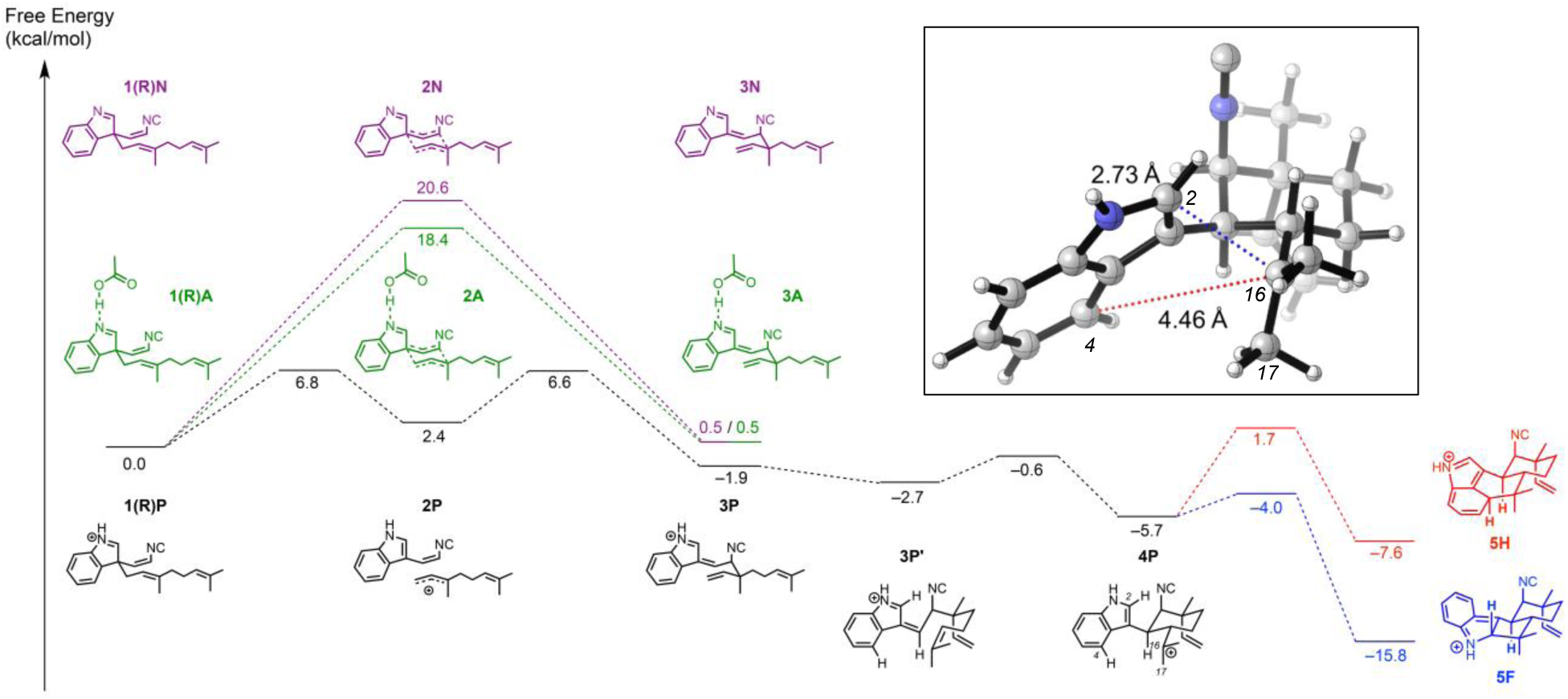
Cope rearrangement, 6-*exo*-*trig* cyclization, and electrophilic aromatic substitution cascade starting from the (*R*)-enantiomer of substrate **1** in a near attack conformation and leading to **12H** precursor **5H** and **12F** precursor **5F**. The energetics of the Cope rearrangement are computed with the neutral indole (pathway **N**), the *N*-protonated indole (pathway **P**), and the indole forming a hydrogen bond with acetic acid (pathway **A**). Inset: Optimized geometry of key intermediate **4P** that undergoes regioselective electrophilic aromatic substitution to form **12H** or **12F**.

The first proposed step in the biosynthesis of **12H** and **12F** is the Cope rearrangement of **1** to intermediate **3**. Although a typical Cope rearrangement has four possible transition states, two chairs and two boats (Fig. S17-S18), only one chair-like transition state can account for the known stereochemistry at positions 11 and 12 in the final products and, therefore, only this transition state was further considered. Given that the otherwise hydrophobic active site of HpiC1 contains the above-described essential D214 residue, we explored the impact of this aspartic acid residue on accelerating the Cope rearrangement. The conversion of neutral starting material **1**(*R*)**N** to intermediate **3N** is concerted, proceeding through a single chair-like transition state **2N**, which lies 20.6 kcal/mol above the near attack conformation (NAC) of the starting material. This chair-like transition structure (Fig. S19) has dissociative character, with breaking and forming partial single bonds of 2.58 Å and 2.53 Å, respectively. The partial negative charge on the isocyanidovinylindoline fragment of –0.43 e can be stabilized by a hydrogen bonding donor at the indole nitrogen, which has a partial negative charge of –0.49 e. Adding an acetic acid molecule to mimic possible hydrogen-bonding between D214 and the indole nitrogen lowers this barrier by 2.2 kcal/mol (40-fold rate enhancement), with the conversion of **1**(*R*)**A** to **3A** proceeding through a single chair-like transition state **2A** with a free energy barrier of 18.4 kcal/mol. These computations suggest that D214 could facilitate the Cope rearrangement of **1** to **3** by hydrogen-bonding to the indole nitrogen. Fully protonating the indole nitrogen, which represents the maximum limit of potential acid catalysis by D214, results in a change of mechanism. The conversion from **1**(*R*)**P** to **3P** is stepwise and dissociative, with **2P** being an intermediate rather than a transition state, and has a much lower overall free energy barrier of 6.8 kcal/mol. Intermediate **2P** is stabilized by full conjugation between the indole and isonitrile groups, and by an allylic cation to produce this much lower overall barrier (Fig. 6).

The second proposed step in the biosynthesis of **12H** and **12F** is the 6-*exo*-*trig* cyclization of intermediate **3** to intermediate **4**, which sets the stereochemistry at positions 10 and 15 in the final products. In gas-phase DFT optimizations, it was only possible to locate a transition state for this cyclization when the indole nitrogen was protonated. Without this protonation, the zwitterionic character of the possible transition state leads to bond formation between negatively-charged position C3 of the indole and positively-charged position C16 to generate a cyclobutane ring. In contrast, the protonated species undergoes facile cyclization from intermediate **3P** to intermediate **4P** through a low-lying transition state. This suggests that protonation is crucial and that D214 may catalyze cyclization in this way, and possibly the preceding Cope rearrangement as well.

Continuing along the protonated pathway, the third step is electrophilic aromatic substitution of intermediate **4P**, with two different transition states leading to the two major products. Electrophilic aromatic substitution at position C4 of the indole yields intermediate **5H**, which gives **12H** upon deprotonation, while electrophilic aromatic substitution at position C2 of the indole yields intermediate **5F**, which gives **12F** upon deprotonation. Rapid deprotonation at the methyl position (C16) instead of electrophilic aromatic substitution can lead to formation of tricyclic **12C** (Fig. 1), which is generated as a trace product by HpiC1 (*23*). QM calculations show that formation of **12F**, which has the lower-energy transition state, should be intrinsically favored. Thus, the regioselectivity of electrophilic aromatic substitution to generate **12H** appears to be controlled by the HpiC1 active site as opposed to inherent energetics of the system.

To obtain additional insights regarding enzymatic control on the initial Cope rearrangement step, we conducted 500 ns MD trajectories on the wild-type HpiC1 with both **1**(*R*) and **1**(*S*) substrate enantiomers as their protonated states, **1**(*R*)**P** and **1**(*S*)**P**, respectively, bound into the enzyme active site (Fig. S20). These MD simulations revealed that both enantiomers retain the key H-bond interaction between the protonated indole NH and the D214 residue for the entirety of the MD trajectories (Fig. S20). However, the C11-C12 distance, which corresponds to the new C-C bond formed during the Cope rearrangement, is shorter (~3.5Å) for the **1**(*R*)**P** substrate than for the **1**(*S*)**P** substrate (>4.0Å). In addition, substrate **1**(*R*)**P** is stabilized by the enzyme active site in a near attack conformation (NAC) that leads to the correct stereochemistry at positions 11 and 12, (*R*) and (*S*), respectively (Fig. 5*c*-*d*). In contrast, substrate **1**(*S*)**P** never explores a NAC that could lead to the correct stereochemistry at positions 11 and 12 observed in the **12H** natural product (Fig. S20-S21). Based on these observations, the (*R*) enantiomer **1**(*R*)**P** is the most plausible natural substrate for the HpiC1 enzyme.

To further understand how the HpiC1 active site could control the formation of the hapalindole/fischerindole products, and how select mutations can alter the product distribution, we performed additional MD simulations with intermediates prior to electrophilic aromatic substitution bound into the active site of the HpiC1 enzyme and associated mutants. We analyzed the impact of the active site in dictating the regioselectivity of the reaction (to form hapalindole or fischerindole cores) by considering the two intermediate precursors **4P** and **8P**, characterized during our QM modeling, that derive from the **1**(*R*)**P** and **1**(*S*)**P** starting materials, respectively. Since either intermediates **4P** or **8P** can generate both the final **12H** and **12F** products, we analyzed the binding of both intermediates in the HpiC1 active site. MD simulations with the two docked intermediates show that intermediate **4P**, derived from the **1**(*R*)**P** substrate, is more effectively bound into the active site, maintaining the H-bond between the protonated NH-indole and D214 more consistently than the **8P** intermediate, which rapidly dissociates during the simulation (Fig. S22). These results reinforce the idea that the **1**(*R*)**P** substrate is the most plausible natural substrate for the HpiC1 enzyme. Moreover, MD simulations showed that when the **4P** intermediate is bound into the active site, it adopts a conformation in which the distance between C4 and C16 (~3.5Å) that corresponds to hapalindole product formation is shorter than the distance between C2 and C16 (~4.0Å) that corresponds to fischerindole product formation (Fig. 5*e*, Fig. S22). Although the intrinsic fischerindole-forming TS is lower in energy than the hapalindole-forming TS, the enzyme active site promotes the formation of the intrinsically less-favored hapalindole product. Finally, late during the MD simulation (around 200 ns) a conformational change of the F138 side chain occurs, which reverses this trend to finally disrupt the interaction of **4P** and D214 (Fig. S22-S23), highlighting the key role of F138 residue in controlling the enzyme activity and also the site-selectivity of the reaction.

The critical role of F138 has been investigated through further MD simulations on the F138S and Y101F/F138S mutants with **4P** intermediate bound. In both trajectories, we observed that the distance between C2 and C16 leading to fischerindole product formation is shortened compared to the wild-type enzyme, becoming similar and slightly closer than the distance between C4 and C16 leading to hapalindole product formation, consistent with the experimentally observed product ratios (Fig. 5*f*, Fig. S24). The absence of the bulky F138 residue near the C2 position allows the intermediate to adopt a slightly different conformation when it is interacting with the D214 residue (Fig. 5*f*, Fig. S25), enabling **12F** formation without completely suppressing generation of **12H**.

## DISCUSSION

The structure of HpiC1 has provided the first high resolution insights into a fascinating mechanistic puzzle in which the cyanobacterial indole alkaloid Stig cyclases are able to generate extensive stereochemical and regiochemical diversity through the central biosynthetic precursor **1**. The surprising function of the Stig cyclases (*10*) could not be inferred from bioinformatic analysis and annotation, and similarly, structural and mechanistic insights from homology-based tools were unavailable for these remarkable biocatalysts. HpiC1 does not share structural homology with any characterized terpene cyclase, but instead shares highest structural homology with bacterial carbohydrate binding modules. CBMs function primarily to bring various hydrolytic enzymes into contact with their carbohydrate substrates. These sugar molecules bind the CBMs in an extended cleft at the protein surface, which is mediated through several amino acids that are not conserved in HpiC1 (Fig. 2*c*) (*20*). This indicates a divergent functionality in HpiC1 that is based on a shared protein scaffold.

Several biochemical characteristics of the CBMs appear to be shared in the Stig cyclases. For instance, thermostability has been reported for CBMs (*24*), and this characteristic was highly beneficial for facile purification of an HpiC1 homolog, FamC1, from the *Fischerella ambigua* cell-free lysate (*10*). As observed with other thermophilic proteins (*25*) an extended salt bridge array in HpiC1 is readily apparent, and spans the dimeric interface in coordination with hydrogen bonds (Fig. S26). These residues are conserved in other Stig cyclases and we expect that many will share this thermostable property. Another feature that is shared between HpiC1 and CBMs is the presence of structural calcium ions. This co-factor also plays an important role in CBM stabilization, substrate recognition, and oligomerization (*24*, *26*).

The molecular basis for the Ca^2+^ dependence of HpiC1 was demonstrated through two well-ordered binding sites in the enzyme. A paradoxical aspect of this calcium requirement is the observation that low millimolar concentrations of CaCl_2_ cause HpiC1 to precipitate. This effect is reversible with stoichiometric addition of EDTA, suggesting a possible calcium-dependent higher order oligomerization of HpiC1. Because HpiC1 could not be crystallized without supplemental CaCl_2_, we examined the crystal packing interactions for any evidence of additional calcium binding sites in the protein. We observed one clearly-resolved interfacial calcium ion (Fig. S27), and a second Ca^2+^ at half occupancy in the native *P*4_2_ crystal. This second bridging calcium ion was also observed in the SeMet crystal despite its unrelated spacegroup, *P*2_1_2_1_2_1_ (Fig. S28), which suggests that the bridging calcium ions could play a role in higher order complexes of the protein. The possible functional role of higher order oligomerization in HpiC1 remains unclear as the addition of supplemental (1 – 20 mM) Ca^2+^ enhances the enzymatic activity of HpiC1, but is not required for turnover (Fig. 4). With some cyclases, a requirement for millimolar concentrations of Ca^2+^ in the reaction buffer to achieve activity has been reported (*15*). Taken together these data are indicative of an important, yet complex structural role for Ca^2+^ in the Stig cyclases. As noted recently and confirmed in the HpiC1 structure, Stig cyclase activity is calcium dependent (*15*), particularly in the fascinating instances of heteromeric associations of the Stig cyclase proteins, leading to variant stereochemical outcomes of the products compared to homodimeric combinations of the biocatalysts (*14*, *15*). Further structural investigation will be required to understand the assembly of heteromeric Stig cyclase complexes, and the relevance of Ca^2+^ in those cases.

We have established the location of the HpiC1 enzyme active site, and using mutational analysis identified critical residues for catalysis and demonstrated a key relationship between amino acid sequence and product outcome. Most importantly, we identified D214 as the source of an active site acid that is required for catalysis, and is consistent with numerous reports of acid catalyzed Cope rearrangements (*22*). Notably, the hydrophobic environment around D214 is essential to maintain a suitable population of the protonated species, as the Y101S/F138S mutation significantly reduced enzyme activity. We also identified a key regiochemical switch at the F138 position that gave rise to production of the fischerindole core in HpiC1. We also confirmed the general importance of this amino acid in the HpiC1 homolog FimC5, where a corresponding mutation also affected the product distribution with respect to regiochemistry (Fig. S8). Together these findings will facilitate our efforts to anticipate the product profiles in both uncharacterized cyclases and new ones that are uncovered as additional strains and gene clusters are discovered through sequencing of relevant microbial genomes.

Using a combination of DFT calculations and MD simulations we have explored the HpiC1 active site dynamics, the origins of its three-part catalytic mechanism, and how HpiC1 controls the regiochemistry of product formation by favoring a particular conformation of substrate **1** and the reaction intermediate **4P**. We have also examined the role of key mutations in HpiC1 that switch the native product outcome from hapalindole **12H** to fischerindole **12F**. Together, these results address several of the catalytic steps in the formation of **12H** from **1**(*R*), although further analysis will be required to establish the mechanistic basis by which the variant Stig cyclases achieve differentiation at the 6-*exo*-*trig* cyclization step, when the stereocenters at C10 and C15 are set (e.g. Hapalindole U, H, and J series) (Fig. S1). The structural and computational analysis presented in this work provide the groundwork to address these questions. And, as demonstrated here, structural studies on additional Stig cyclases, mutational analysis across key active site residues, and computational modeling of the reaction intermediates will enable greater ability to predict product profiles, and engineer new selectivities to diversify further this remarkable family of natural products.

## EXPERIMENTAL DESIGN

### Cloning and mutagenesis of HpiC1 and FimC5

HpiC1 and FimC5 were cloned into pET28 (Novagen) from codon optimized synthetic genes (IDT gBlocks^®^) with their N-terminal leader peptides truncated (*14*). Site-directed mutagenesis of HpiC1 and FimC5 was performed using a single primer method based on “Quikchange” mutagenesis (Agilent Genomics). Mutagenic primer sequences are listed in Table S1. All mutations were verified by DNA sequencing at the University of Michigan DNA Sequencing Core.

### Expression of HpiC1 and FimC5 proteins

HpiC1 and FimC5 and their corresponding active site mutants were overexpressed in *Escherichia coli* strain BL21 (DE3). Cultures from a single colony were used to inoculate 1.5 L terrific broth (TB) supplemented with 50 μg/mL kanamycin. Expression was induced with 0.7 mM isopropyl-β-D-thiogalactopyranoside when cultures reached OD_600_ ~ 1.0. After 20 h induction at 18 °C the cells were harvested by centrifugation and stored at −80 °C.

### Expression of HpiC1 W73M/K132M selenomethionine derivative

An initial challenge involved the lack of native methionine residues in HpiC1. Therefore, a series of mutants containing methionine substitutions were screened for crystallization. Sites for substitution were selected based on positions containing a native methionine in sequence comparisons with other Stig cyclases. Selenomethionine (SeMet) HpiC1 W73M/K132M was produced by metabolic inhibition (*27*). Briefly, freshly transformed BL21 (DE3) cells harboring the hpiC1 gene on pET28 were used to incolulate 3 L M9 minimal medium supplemented with 50 μg/mL kanamycin. An amino acid cocktail containing L-selenomethionine was added when the cells reached OD_600_ = 1.0. The cells were cooled to 18 °C and shaken for 30 min prior to induction with 0.7 mM isopropyl-β-D-thiogalactopyranoside. After 20 h induction the cells were harvested by centrifugation and stored at −80 °C.

### Purification of recombinant proteins

All proteins were purified as described previously (*14*). Briefly, 10 g *E. coli* wet cell mass containing the recombinant cyclase was resuspended in 75 mL lysis buffer (10 mM HEPES pH 7.6, 50 mM NaCl, 10% glycerol). Cells were lysed by the addition of lysozyme (0.5 mg/mL) and sonication and clarified by centrifugation at 60,000 x*g* for 25 min. Clarified lysate was loaded by gravity onto 8 mL NiNTA Superflow resin (Qiagen). The column was washed with 100 mL lysis buffer containing 20 mM imidazole and 50 mL lysis buffer containing 40 mM imidazole. The proteins were eluted with elution buffer (250 mM imidazole, pH 7.9 and 10% glycerol). Fractions containing the purified cyclase were concentrated using Amicon Ultra 15 centrifugal filters and desalted using PD-10 columns (GE Healthcare) equilibrated with storage buffer (10 mM HEPES pH 7.6, 50 mM NaCl). The purified cyclases were drop-frozen in 30 μL aliquots directly into liquid N_2_ and stored at −80 °C.

### Crystallization of SeMet HpiC1 W73M/K132M

Single, diffraction quality crystals of the HpiC1 W73M/K132M selenomethionine derivative were grown in Intelli-Plate 96-2 shallow well plates (Hampton research) at 20 °C by mixing 1 μL of 11 mg/mL SeMet HpiC1 in storage buffer with 1 μL of a well solution containing 23% PEG 3350, 200 mM CaCl_2_, 5% trehalose. Sitting droplets were nucleated after 18 h from an earlier spontaneous crystallization event using a cat whisker. Single, rod-shaped crystals grew to approximate dimensions of 50 x 50 x 250 μm after 14 days. 8 μL of a cryoprotecting solution containing 10 mM HEPES pH 7.6, 50 mM NaCl, 23% PEG 3350, 200 mM CaCl_2_, 9.1% trehalose was added directly to the sitting drops and the crystals were harvested using nylon loops and vitrified by rapid plunging into liquid nitrogen. SeMet HpiC1 crystallized in Form 1, space group *P*2_1_2_1_2_1_ with unit cell dimensions of *a* = 44.9 Å, *b* = 81.1 Å, *c* = 131.7 Å, and two chains in the asymmetric unit. *Crystallization of Native HpiC1 (P4_2_)*. Single, diffraction quality crystals of HpiC1 native were grown in Intelli-Plate 96-2 shallow well plates (Hampton research) at 20 °C by mixing 1 μL of 20 mg/mL HpiC1 in storage buffer with 1 μL of a well solution containing 22% PEG 4000, 200 mM CaCl_2_, 100 mM Tris pH 8.5, 5% ethylene glycol. Sitting droplets were nucleated after 4 h from an earlier spontaneous crystallization event using a cat whisker. Single, rod-shaped crystals grew to approximate dimensions of 50 x 50 x 150 μm after 7 days. 8 μL of a cryoprotecting solution containing 10 mM HEPES pH 7.6, 50 mM NaCl, 22% PEG 4000, 200 mM CaCl_2_, 100 mM Tris pH 8.5, 15% ethylene glycol was added directly to the sitting drops and the crystals were harvested using nylon loops and vitrified by rapid plunging into liquid nitrogen. In these conditions, HpiC1 native crystallized in Form 2, space group *P*4_2_ with unit cell dimensions of *a* = 71.3 Å, *b* = 71.3 Å, *c* = 80.6 Å, and two chains in the asymmetric unit.

### Crystallization of Native HpiC1 (C2)

Single, diffraction quality crystals of HpiC1 native were grown in Intelli-Plate 96-2 shallow well plates (Hampton research) at 20 °C by mixing 1 μL of 20 mg/mL HpiC1 in storage buffer and 5% DMSO with 1 μL of a well solution containing 22% PEG 4000, 150 mM CaCl_2_, 100 mM Tris pH 8.5, 5% ethylene glycol. Sitting droplets were nucleated after 4 h from an earlier spontaneous crystallization event using a cat whisker. Single, diamond-shaped crystals grew to approximate dimensions of 200 x 200 x 100 μm after 7 days. 8 μL of a cryoprotecting solution containing 10 mM HEPES pH 7.6, 50 mM NaCl, 22% PEG 4000, 150 mM CaCl_2_, 100 mM Tris pH 8.5, 15% ethylene glycol, 5% DMSO was added directly to the sitting drops and the crystals were harvested using nylon loops and vitrified by rapid plunging into liquid nitrogen. In these conditions, HpiC1 native crystallized in Form 3, space group C2 with unit cell dimensions of *a* = 113.8 Å, *b* = 49.5 Å, *c* = 53.1 Å, α = 90°, β = 110.5°, γ = 90° and one chain in the asymmetric unit.

### Crystallization of HpiC1 Y101F

Single, diffraction quality crystals of HpiC1 Y101F were grown in Intelli-Plate 96-2 shallow well plates (Hampton research) at 20 °C by mixing 1 μL of 15 mg/mL protein in storage buffer with 1 μL of a well solution containing 22% PEG 4000, 150 mM CaCl_2_, 100 mM Tris pH 8.5, 5% ethylene glycol. Sitting droplets were nucleated after 4 h from an earlier spontaneous crystallization event using a cat whisker. Single, diamond-shaped crystals grew to approximate dimensions of 250 x 250 x 270 μm after 7 days. 8 μL of a cryoprotecting solution containing 10 mM HEPES pH 7.6, 50 mM NaCl, 22% PEG 4000, 150 mM CaCfe, 100 mM Tris pH 8.5, 15% ethylene glycol was added directly to the sitting drops and the crystals were harvested using nylon loops and vitrified by rapid plunging into liquid nitrogen. HpiC1 Y101F crystallized in Form 3, space group *C*2 with unit cell dimensions of *a* = 113.8 Å, *b* = 49.8 Å, *c* = 53.4 Å, α = 90°, β = 110.4°, γ = 90° and one chain in the asymmetric unit.

### Crystallization of HpiC1 Y101S

Single, diffraction quality crystals of HpiC1 Y101S were grown in Intelli-Plate 96-2 shallow well plates (Hampton research) at 20 °C by mixing 1 μL of 15 mg/mL protein in storage buffer with 1 μL of a well solution containing 20% MEPEG 5000, 150 mM CaCl_2_, 100 mM Tris pH 8.5, 5% ethylene glycol. Sitting droplets were nucleated after 4 h from an earlier spontaneous crystallization event using a cat whisker. Single, diamond-shaped crystals grew to approximate dimensions of 250 x 250 x 270 μm after 7 days. 8 μL of a cryoprotecting solution containing 10 mM HEPES pH 7.6, 50 mM NaCl, 20% MEPEG 5000, 150 mM CaCl_2_, 100 mM Tris pH 8.5, 15% ethylene glycol, 5% DMSO was added directly to the sitting drops and the crystals were harvested using nylon loops and vitrified by rapid plunging into liquid nitrogen. HpiC1 Y101S crystallized in Form 3, space group C2 with unit cell dimensions of *a* = 113.9 Å, *b* = 49.6 Å, *c* = 53.4 Å, α = 90°, β = 110.3°, γ = 90° and one chain in the asymmetric unit.

### Crystallization of HpiC1 F138S and Y101F/F138S

Single, diffraction quality crystals of HpiC1 F138S and Y101F/F138S were grown in Intelli-Plate 96-2 shallow well plates (Hampton research) at 20 °C by mixing 1 μL of 15 mg/mL protein in storage buffer, 20 mM CaCl_2_, 5% DMSO with 1 μL of a well solution containing 20% MEPEG 5000, 100 mM BisTris pH 6.5, 5% ethylene glycol. Sitting droplets were nucleated after 4 h from an earlier spontaneous crystallization event using a cat whisker. Single, plate-shaped crystals grew to approximate dimensions of 50 x 50 x 300 μm after 7 days. 8 μL of a cryoprotecting solution containing 10 mM HEPES pH 7.6, 50 mM NaCl, 20% MEPEG 5000, 20 mM CaCl_2_, 100 mM BisTris pH 6.5, 15% ethylene glycol, 5% DMSO was added directly to the sitting drops and the crystals were harvested using nylon loops and vitrified by rapid plunging into liquid nitrogen. HpiC1 F138S and HpiC1 Y101F/F138S crystallized in Form 4, space group *P*2_1_ with unit cell dimensions of *a* = 62.0 Å, *b* = 47.9 Å, *c* = 174.2 Å, α = 90°, β = 97.2°, γ = 90° and four chains in the asymmetric unit. *Data collection and processing.* X-ray data were collected at 100 K on beamline 23ID-B at the General Medical Sciences and Cancer Institutes Structural Biology Facility at the Advanced Photon Source in Argonne, IL, USA. Diffraction data were integrated and scaled using XDS (*28*). Data collection statistics are given in Table 1.

### Experimental phasing (SAD) and molecular replacement, model building and refinement

The structure of SeMet HpiC1 W73M/K132M was solved using single wavelength anomalous diffraction (SAD). Phasing and initial model building were performed using Phenix Autosol (*29*). This resulted in an initial model that could be extended by alternating cycles of manual building in *Coot* (*30*) and least-squares refinement with Refmac (*31*). The structures for HpiC1 native and Y101F, Y101S, F138S, Y101F/F138S were solved by molecular replacement using Phaser-MR (*32*) with the structure of the HpiC1 selenomethionine derivative as a search model. Final models were generated by alternating cycles of manual building in *Coot* (*30*) and refinement in Refmac (*31*) and Phenix (*33*), and were validated using MolProbity (*34*). *Docking 12*-*epi*-*hapalindole U with Autodock VINA*. 12-*epi*-hapalindole U was docked into the SeMet HpiC1 model using Autodock VINA (*21*). Default parameters for Autodock VINA were used with the exception of exhaustiveness, which was set to 100.

### Chemical synthesis

Indole isonitrile was synthesized as described previously (*10*).

### In vitro cyclase assays

In vitro assays were performed with HpiC1, FimC5 and their corresponding active site mutants as described previously (*14*). Briefly, FamD2, GPP, Indole isonitrile, cyclase, CaCl_2_. Products were analyzed using LC/MS (Shimadzu) using C18 (Agilent) HPLC column.

### Scaleup, purification, and NMR of F138Sproduct (12-epi-fischerindole U)

The semi-prep scale reaction was performed as described previously (*14*).

### Quantum mechanical calculations

Conformational searches of the hapalindole and fischerindole products were performed using the Schrödinger MacroModel (*35*) software package, and the lowest energy conformation was used for all reported quantum mechanical calculations. All quantum mechanical calculations were performed using the Gaussian 09 (*36*) software package. Structures were optimized in the gas phase at the B3LYP (*37*, *38*)/6-31G(d) level of theory; frequency calculations were used to confirm the presence of local minima (no imaginary frequencies) and transition states (one imaginary frequency) and to calculate free energies at 298 K. To obtain more accurate energetics, single-point energy calculations were performed on the optimized structures at the B3LYP/6-311++G(d,p) level of theory using Grimme’s D3(BJ) dispersion correction (*39*, *40*) and the IEFPCM (*41*-*43*) solvent model for diethyl ether (ε = 4). The use of the dielectric constant ε=4 has proven to be a good model to estimate the dielectric permittivity in the enzyme active site, accounting for electronic polarization and small backbone fluctuations (*44*, *45*).

### Molecular Dynamics simulations

Molecular Dynamics simulations were performed using the GPU code (*pmemd*) (*46*) of the AMBER 16 package (*47*). Parameters for intermediates and substrates were generated within the *antechamber* module using the general AMBER force field (*gaff*) (*48*), with partial charges set to fit the electrostatic potential generated at the HF/6-31G(d) level by the RESP model (*49*). The charges were calculated according to the Merz–Singh–Kollman scheme (*50*, *51*) using the Gaussian 09 package (*36*). Each protein was immersed in a pre-equilibrated truncated cuboid box with a 10 Å buffer of TIP3P (*52*) water molecules using the *leap* module, resulting in the addition of around 15,000 solvent molecules. The systems were neutralized by addition of explicit counter ions (Na^+^ and Cl^−^). All subsequent calculations were done using the widely tested Stony Brook modification of the Amber14 force field (*ff14sb*) (*53*). A two-stage geometry optimization approach was performed. The first stage minimizes the positions of solvent molecules and ions imposing positional restraints on the solute by a harmonic potential with a force constant of 500 kcal·mol^−1^·Å^−2^ and the second stage minimizes all the atoms in the simulation cell. The systems were gently heated using six 50 ps steps, incrementing the temperature by 50 K for each step (0–300 K) under constant-volume and periodic-boundary conditions. Water molecules were treated with the SHAKE algorithm such that the angle between the hydrogen atoms was kept fixed. Long-range electrostatic effects were modelled using the particle-mesh-Ewald method (*54*). An 8 Å cutoff was applied to Lennard–Jones and electrostatic interactions. Harmonic restraints of 10 kcal·mol^−1^ were applied to the solute and the Langevin equilibration scheme was used to control and equalize the temperature. The time step was kept at 1 fs during the heating stages, allowing potential inhomogeneities to self-adjust. Each system was then equilibrated without restrains for 2 ns with a 2 fs time step at a constant pressure of 1 atm and temperature of 300 K. After the systems were equilibrated in the NPT ensemble, subsequent MD simulations were performed for an additional 500 ns under an NVT ensemble and periodic-boundary conditions.

